# LPS induces GFAT2 expression to promote O-GlcNAcylation and attenuate inflammation in macrophages

**DOI:** 10.1101/2020.03.22.002303

**Authors:** Hasanain Al-Mukh, Léa Baudoin, Abdelouahab Bouaboud, José-Luis Sanchez-Salgado, Nabih Maraqa, Mostafa Khair, Patrick Pagesy, Georges Bismuth, Florence Niedergang, Tarik Issad

## Abstract

O-GlcNAc glycosylation is a reversible post-translational modification that regulates the activity of intracellular proteins according to glucose availability and its metabolism through the hexosamine biosynthesis pathway (HBP). This modification has been involved in the regulation of various immune cell types, including macrophages. However, little is known concerning the mechanisms that regulate protein O-GlcNAcylation level in these cells. In the present work, we demonstrate that LPS treatment induces a marked increase in protein O-GlcNAcylation in RAW264.7 cells, bone-marrow-derived and peritoneal mouse macrophages, as well as human monocyte-derived macrophages. Targeted deletion of OGT in macrophages resulted in an increased effect of LPS on NOS2 expression and cytokine production, suggesting that O-GlcNAcylation may restrain inflammatory processes induced by TLR4 activation. The effect of LPS on protein O-GlcNAcylation in macrophages was associated with an increased expression and activity of glutamine fructose 6-phosphate amido-transferase (GFAT), the enzyme that catalyzes the rate-limiting step of the HBP. More specifically, we observed that LPS rapidly and potently stimulated GFAT2 isoform mRNA and protein expression. Genetic or pharmacological inhibition of FoxO1 impaired LPS effect on GFAT2 expression, suggesting a FoxO1-dependent mechanism. We conclude that *GFAT2* should be considered as a new TLR4-inducible gene involved in regulation of protein O-GlcNAcylation, that permits to limit exacerbated inflammation upon macrophage activation.

## Introduction

O-GlcNAcylation is a post-translational modification that regulates the activity of cytosolic, nuclear and mitochondrial proteins. This modification is controlled by only two enzymes: OGT, that adds N-Acetylglucosamine (GlcNAc) on serine or threonine residues, and OGA, that removes it. O-GlcNAcylation, which regulates a wide array of biological processes [1], depends on the availability of glucose and its metabolism through the hexosamine biosynthesis pathway (**Suppl. Fig S1**). This modification has been involved in the modulation of various signalling pathways, and perturbations of O-GlcNAcylation participate in several important human pathologies [2-4]. Moreover, a number of studies indicated a link between O-GlcNAcylation and inflammatory conditions, including diabetes, auto-immune diseases and cancer [5-11]. Moreover, several lines of evidence have suggested a role for O-GlcNAcylation in immune cell signalling. Indeed, O-GlcNAcylation was first discovered at the surface of lymphocytes [12] and dynamic changes upon lymphocyte activation were described at the beginning of the nineties [13]. More recently, several studies indicated that protein O-GlcNAcylation plays major roles in the immune system. In T cells, TCR (T-cell receptor) activation results in global elevation of protein O-GlcNAcylation [14,15], increased O-GlcNAcylation of several transcription factors [16-19] and accumulation of OGT at the immunological synapse [14]. Moreover, in B lymphocytes, activation of the B-cell receptor (BCR) induces O-GlcNAcylation of several signalling molecules involved in BCR signalling [20,21]. Overall, these studies strongly support a major role in O-GlcNAcylation in adaptive immunity. However, the molecular mechanisms by which immune cell activation modulates protein O-GlcNAcylation remain elusive.

In macrophages, contradictory results have been obtained concerning the effect of LPS on protein O-GlcNAcylation and its role in inflammatory processes. Whereas some studies indicated that LPS induced a general increase in protein O-GlcNAcylation [22,23], others found that LPS decreases protein O-GlcNAcylation in macrophages [24,25]. Moreover, whereas several lines of evidence indicate that increased O-GlcNAcylation of signaling proteins in the NFκB pathway potentiates inflammatory processes [26-29], other studies indicate that increased O-GlcNAcylation may inhibit pro-inflammatory signals [24,30]. More recently, O-GlcNAcylation of the signalling adaptor MAVS in macrophages has been involved in antiviral signalling response [31], whereas increased RIPK3 O-GlcNAcylation was shown to regulate necroptosis signalling upon LPS stimulation [25]. Thus, whereas numerous evidences suggest that modulation of O-GlcNAcylation signalling constitutes an important facet of innate immune response, the mechanisms by which macrophage activation regulates O-GlcNAc, and its consequences on inflammatory processes, remain to be firmly established.

In the present work, using different macrophage cell models, we clearly demonstrated that TLR4 activation markedly stimulates protein O-GlcNAcylation in macrophages through increased expression and activity of the rate-limiting enzyme of the hexosamine biosynthetic pathway, glutamine fructose-6-phosphate amidotransferase (GFAT). More specifically, we showed that whereas resting macrophages express mainly the GFAT1 isoform, upon LPS stimulation, a marked increase in GFAT2 expression was observed, revealing GFAT2 as a new TLR4-inducible gene in macrophages that promotes a general increase in protein O-GlcNAcylation. In macrophages with conditional OGT deletion, lack of LPS-induced O-GlcNAyclation was associated with an increased in the production of pro-inflammatory molecules, suggesting that GFAT2 induction may be part of a regulatory loop that may limit inflammation and/or permit its resolution after TLR4 activation.

## Materiel and Methods

### Chemicals and antibodies

Thiamet G (3aR,5R,6S,7R,7aR)-2-(ethylamino)-3a,6,7,7a-tetrahydro-5-(hydroxymethyl)-5H-Pyrano.thiazole-6,7-diol) and LPS (Lipopolysaccharides) from Salmonella enterica serotype typhimurium were from Sigma-Aldrich (Saint Quentin Fallavier France). FoxO1 inhibitor AS1842856 was from Calbiochem. PUGNAc (*O*-(2-acetamido*-*2-deoxy-d-glucopyranosylidene)-amino*-N*-phenylcarbamate) was from Toronto Research Chemicals, Inc. (North York, ON, Canada). A list of antibodies used in this study is given in Supplementary Table 1.

### cDNA constructs

The cDNA coding for nuclear, cytosolic and plasma membrane FRET O-GlcNAc biosensors [32] were generously provided by Prof. L.K. Mahal (University of Texas, Austin, USA). BRET biosensors were developed based on these FRET constructs by replacing the CFP by a Rluc8 sequence [33].

Human (−801>>>0) and mouse GFAT2 (−501>>>0) putative promoter sequences were amplified by PCR and cloned in a firefly luciferase plasmid (pGL4-20, Promega). FoxO1 binding sites on GFAT2 promoter were identified using Regulatory Sequence Analysis Tools (RSAT) web server (http://rsat.ulb.ac.be/rsat/). Mutagenesis of Foxo1 binding sites on the mouse GFAT2 reporter gene was performed using mutagenesis kit (QuikChangeII XL-Agilent Technologies).

Constitutively active FOXO1-TM (mutated on the 3 Akt phoshorylation sites) has been described previously [34].

### Cell culture and transfection

RAW264.7 murine macrophage cells were maintained in media constituted of RPMI 1640-glutamax medium supplemented with 10% fetal calf serum (FCS), 50µM β-mercaptoethanol, 1 mM sodium pyruvate, 10 mM HEPES and 2 mM L-Glutamine (GIBCO).

Plasmid transfections were performed by cell electroporation using the Electrobuffer kit (Cell projects Ltd). For each transfection, cells grown to sub-confluence in a 100 mm plate were transfected with 15 µg of plasmid DNA. Cells were electroporated at 250 V, 900 µF in 0.4 cm cuvettes (Biorad) using the Gene Pulser X electroporation system (Bio-Rad Laboratories). After electroporation, cells were immediately re-suspended in culture medium, transferred into 96-well White Optiplates previoulsy coated with polylysine (Perkin Elmer) and then cultured for 18 hours at 37°C and 5% CO2 before treatments.

HEK 293-T cells were cultured in DMEM and transfected with Fugene as described previously [34].

### Chromatin immunoprecipitation (ChIP)

RAW264.7 cells (1×10^7^ cells) were prepared for ChIP assay using HighCell# ChIP kit protein A (Diagenode). After cross-linking with formaldehyde, DNA was sonicated into 200-300bp fragments, and protein-DNA complexes were immunoprecipitated using either FoxO1-ChIP grade antibody (Abcam, ab39670) or control rabbit IgG. Protein-DNA crosslinks were reversed by heating, and precipitated DNA was quantified by real-time PCR using primers (Table 1) that amplify either Foxo1 binding site 1 or 2 present in the 500bp upstream of the transcription start site of the mouse GFAT2 gene.

### BRET experiments

Transfected cells were treated with LPS (100ng/mL) and/or Thiamet G (10μM) for 24h. BRET experiments were then performed exactly as described previously [35,36] using the Infinite F200 Pro microplate analyser (Tecan). Briefly, cells were pre-incubated for 5 min in PBS in the presence of 5µM coelenterazine. Each measurement corresponded to the signal emitted by the whole population of cells present in a well. BRET signal was expressed in milliBRET Unit (mBU). The BRET unit has been defined previously as the ratio 530 nm/485 nm obtained in cells expressing both luciferase and YFP, corrected by the ratio 530 nm/485 nm obtained under the same experimental conditions in cells expressing only luciferase [37,38].

### Animals

8 to 12 weeks-old C57BL/6J male mice were used. To study the function of Foxo1 in macrophages, we crossed mice carrying two floxed *Foxo1* alleles kindly provided by Prof. Hedricks, Univ. of California, San Diego, USA) with LysM-Cre transgenic mice (kindly provided by Carol Peyssonnaux, Institut Cochin, Paris) in which Cre is specifically expressed in the myelomonocytic cell lineage (LysMCre-Foxo1/Foxo1 knock-out (KO) mice, thereafter refered to as Foxo1-KO mice). To study the role of O-GlcNAcylation on pro-inflammatory effects of LPS, mice with OGT deletion in the myelomonocytic cell lineage were generated by crossing mice carrying two floxed *OGT* alleles (obtained from Jackson laboratories) with LysM-Cre transgenic mice (LysMCre-OGT knock-out (KO) mice, thereafter referred to as OGT-KO mice). All mice were housed in the Institut Cochin animal facility. All experiments were performed in accordance with accepted standards of animal care, as established in the Institut National de la Santé et de la Recherche Médicale (INSERM) and the Centre National de la Recherche Scientifique (CNRS) guidelines and were approved by the national ethical committee (APAFIS N°A751402).

### Preparation of bone-marrow-derived and peritoneal macrophages from mice

Mice were sacrificed by CO_2_ asphyxiation. Bone marrow-derived macrophages (BMDM) were prepared from bone marrow cells flushed from femurs and tibias. Cells were seeded in sterile Petri dishes at the concentration of 10^6^/ml in RPMI medium supplemented with 10% FCS, 100µg/ml gentamycin, 10mM HEPES, 1mM sodium pyruvate, 50µM β-mercaptoethanol and 20ng/mL mouse MCSF (Miltenyi Biotec). Cells were differentiated during 6 days, then washed to eliminate non-adherent cells, and macrophages were detached in cold PBS -/- and seeded in 6 well-plates at 10^6^ cells/ml in complete media. Cells were treated the day after.

For peritoneal macrophages collection, after mice sacrifice, the peritoneum was infused with 10mL of cold RPMI. Cells were seeded in 6 well-plates at 10^6^cells/ml in complete media and allowed to adhere overnight. Non-adherent cells were then washed away and macrophages treated with LPS.

In some experiments, mice were injected intraperitoneally with LPS (0.6 mg/kg). 6h after injection, mice were sacrificed and peritoneal cells were harvested as described above and plated in a 6 well-plate at 37 °C with 5% CO2. After 2 h, non-adherent cells were washed away, and adherent cells were lysed for analysis by western blot.

To evaluate the effect of LPS on cytokine production *in vivo* in OGT-KO mice, LPS was injected intraperitoneally, mice were sacrificed 6 h after injection and blood was collected by cardiac puncture. Blood was centrifuged at 1500 rpm for 10 min at 4°C and the serum was collected and frozen at −80°C for subsequent determination of cytokine concentrations using MSD kit.

### Preparation of Human monocytes derived macrophages

Human primary macrophages were isolated from blood of healthy donors (Etablissement Français du Sang, Ile-de-France, Site Trinité, Agreement number INSERM-EFS:18/EFS/030) by density gradient sedimentation on Ficoll (GE Healthcare), followed by negative selection on magnetic beads (Stem cells, Cat n°19059) and adhesion on plastic at 37°C during 2 h. Cells were then cultured in the presence of complete culture medium (RPMI 1640 supplemented with 10 % FCS (Eurobio), 100 mg/ml streptomycin/penicillin and 2 mM L-glutamine (Gibco) containing 10 ng/ml rhM-CSF (R&D systems) during 4-5 days [39].

### RNA extraction, reverse transcription and qPCR

Macrophages cultured in 6-well plates were lysed in Trizol reagent (Life Technologies). RNA was isolated and reverse transcribed [40]. Levels of the cDNA of interest were measured by qPCR (LightCycler FastStart DNA Master SYBR Green 1 kit; Roche Diagnostics). The absence of genomic DNA contamination was verified by treating RNA samples in parallel without reverse transcriptase and controlling for the absence of amplification by qPCR. Gene expression was normalized over cyclophilin and HPRT (Hypoxanthine-guanine phosphoribosyltransferase) RNA levels. The sequences of the primers used for qPCR are given in Supplementary Table 2.

### OGA enzymatic assay

Control and LPS-treated RAW264.7 cells were lysed in extraction buffer containing 50mM Tris, 137 mM NaCl, 1% triton, 10% glycerol, phosphatases (50 mM NaF, 10 mM di-sodium β-glycerophosphate, 1 mM Na _3_VO_4_) and proteases (AEBSF, leupeptine, antipaine aprotinine and pepstatine, 1μg/ml each) inhibitors. OGA activity was measured using 4-methylumbellifery-N-acetylβ-D-glucosamine (MU-GlcNAc, Sigma), which is converted into fluorescent 4-methylumbelliferon upon hydrolysis by OGA and other hexosaminidases [41]. 4-methylumbelliferon fluorescence was measured at 448 nm after excitation at 362 nm. Fluorescent measurements were performed after 30 min and 60 min of incubation at 37 °C, to ensure that the determination was performed during the linear phase of the reaction. To specifically determine OGA activity versus other hexosaminidase present in the lysate, all reactions were performed both in absence and presence of 100 μM Thiamet G (a specific OGA inhibitor). The difference between the fluorescent signals obtained in absence (activity of OGA plus other hexosaminidases) and presence of Thiamet G (activity of the other hexosaminidases) reflected the amount of 4-methylumbelliferon produced by OGA [9].

### OGT enzymatic assay

Control and LPS-treated RAW264.7 cells were lysed in extraction buffer containing 50mM Tris, 137 mM NaCl, 1% triton, 10% glycerol, phosphatases (50 mM NaF, 10 mM di-sodium β-glycerophosphate, 1 mM Na _3_VO_4_) and proteases (AEBSF, leupeptine, antipaine aprotinine and pepstatine, 1μg/ml each) inhibitors. Immunoprecipitation of OGT was performed as described below. 400 µg of proteins were incubated with 1.5 µg of anti-OGT antibody (Sigma-Aldrich) for 2h at 4°C. Precipitation was performed by incubating 25µL equilibrated protein G-sepharose beads (GE Healthcare) for 30 min at 4°C. After 3 washes, the precipitated proteins were submitted to an additional wash in OGT assay buffer containing 50 mM Tris-HCl and 12.5 mM MgCl_2_, pH7.5. OGT assay was then performed on protein-G sepharose bound OGT [33] using the bioluminescent UDP-Glo™ glycosyltransferase assay (Promega) exactly as described in the manufacturer instructions [42].

### GFAT enzymatic assay

Glutamine fructose-6-phosphate amidotransferase enzymatic activity was measured as described previously [43]. Control and LPS-treated RAW264.7 cells, human monocyte-derived macrophages or mouse bone-marrow-derived macrophages were lysed in extraction buffer containing 50mM Tris, 137 mM NaCl, 1% triton, 10% glycerol, phosphatases (50 mM NaF, 10 mM di-sodium β-glycerophosphate, 1 mM Na _3_VO_4_) and proteases (AEBSF, leupeptine, antipaine aprotinine and pepstatine, 1μg/ml each) inhibitors. 15-30 μg of proteins were incubated in a buffer containing 10 mM fructose 6-phosphate, 6 mM glutamine, 0.3 mM 3-acetylpyridine adenine dinucleotide (APAD), 50 mM KC1, 100 mM KH_2_PO_4_ (pH 7.5), 1 mM dithiothreitol and 30 unit/ml of glutamate dehydrogenase. After incubation for 30 min at 37°C, the change in absorbance due to reduction of APAD to APADH was monitored spectrophotometrically at 365 nm using TECAN™ infinite M1000 Pro Microplate Reader.

### Western blotting

Cells were lysed with buffer containing 50mM Tris–HCl (pH 8), 137mM NaCl, 10% (v/v) glycerol, 1% (v/v) triton, 50mM NaF, 10mM di-sodium β-glycerophosphate, 1mM Na _3_VO_4_, protease inhibitors (1µg/ml pepstatin, antipain, leupeptin, aprotinin and AEBSF), supplemented with 10μM PUGNAc in order to preserve the GlcNAcylation state of proteins during the extraction procedure. Proteins were then analysed by SDS-PAGE followed by western-blotting as described previously [44].

In some experiments, O-GlcNAcylated proteins were precipitated on wheat germ lectin as described previously [45,46]. Briefly, cell lysates (400 μg of proteins) were incubated for 2 hours at 4° C on a rotating wheel with 20 µl of agarose beads coupled to wheat germ lectin (which binds the N-Acetyl-Glucosamine pattern). At the end of the incubation, the beads were pelleted by centrifugation at 3000 g for 2 min. The supernatant was removed and the pellet was washed 3 times in extracting buffer. Twenty µL of Laemmli sample buffer were added to the beads, and the samples were boiled at 95°C for 5 minutes and then submitted to western-blotting as described previously.

### Statistical analysis

Statistical analyses were performed using PRISM software. Comparison between groups were performed using Student’s t test, or ANOVA followed by Dunnett’s or Tukey’s post-test for multiple comparison analysis.

## Results

### LPS stimulation markedly increased protein O-GlcNAcylation in macrophages

Sub-cellular relocalisation of OGT in different cell compartments has been observed upon stimulation of membrane receptors [47,48], resulting in compartment specific changes in O-GlcNAcylation activity. In order to evaluate whether TLR4 stimulation affects protein O-GlcNAcylation in different cellular compartments in macrophages, we used BRET-based O-GlcNAc biosensors [33]. These biosensors are composed of Rluc8 luciferase fused to a lectin domain (GafD), a known OGT substrate peptide derived from casein kinase II, followed by the Venus variant of the yellow fluorescent protein (**Fig. 1A**). Upon O-GlcNAcylation, the casein kinase peptide binds to the lectin, resulting into a conformational change detected as an increased BRET signal. These biosensors were fused to addressing sequences for targeting to the internal face of the plasma membrane (using Lyn myristoylation/palmitoylation sequence), the cytosol (using the HIV-1 Rev protein nuclear exclusion sequence) or the nucleus (using the SV40 nuclear localisation sequence) [32].

**Figure 1:**
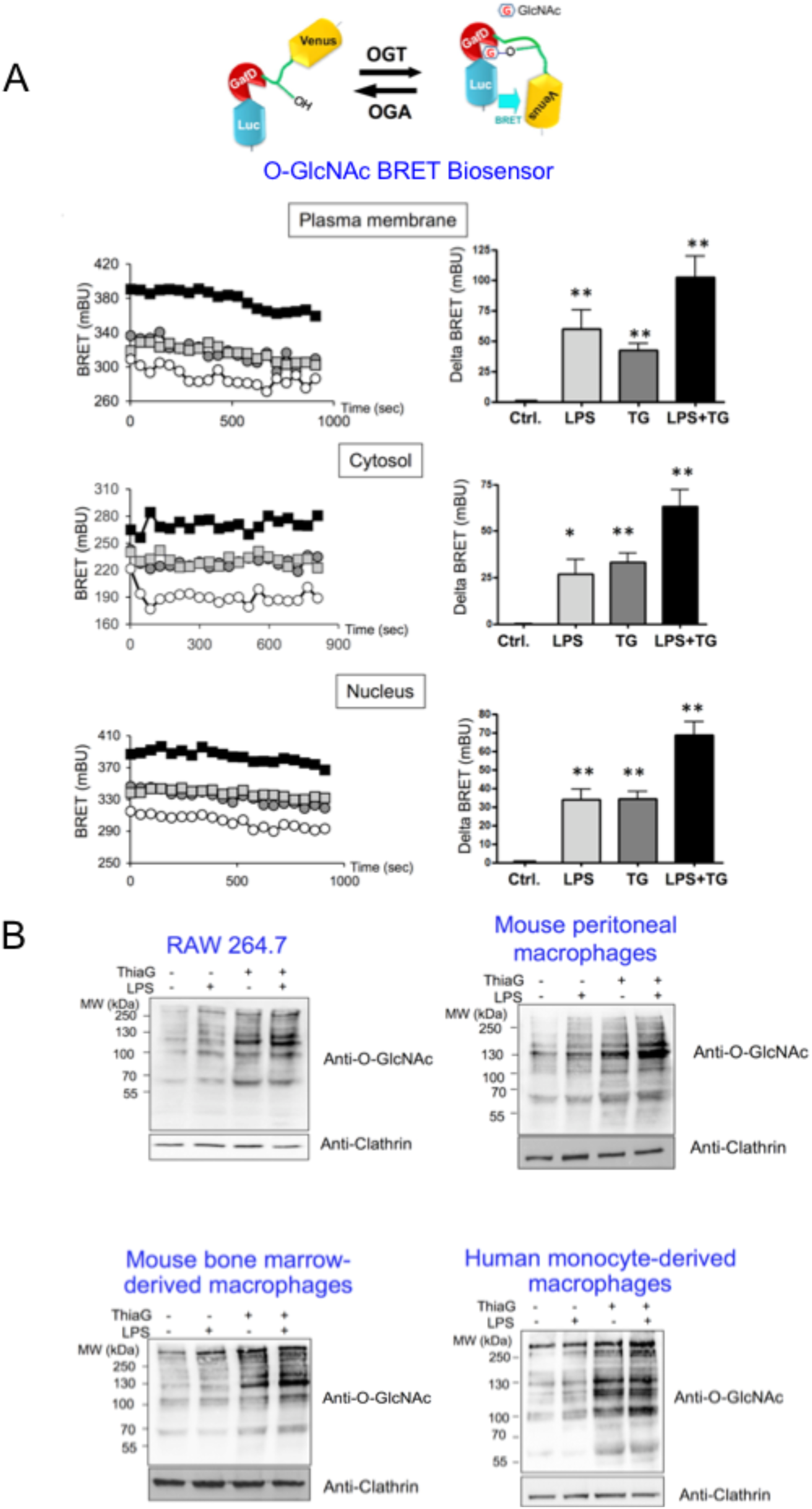
LPS induces O-GlcNAcylation in macrophages. (A) RAW264.7 cells were transfected with plasma membrane, cytosol or nucleus-targeted BRET biosensors. 18h after transfection, medium was changed and cells were cultured for an additional 24h in presence of LPS (100ng/mL), Thiamet G (TG, 10µM) or both. BRET signal was then measured every 45 seconds during 15 min. In each experiment, the mean of 20 repeated BRET measurements in a given experimental condition was taken as the BRET value obtained in this experimental condition. Left panels show the signals obtained during typical BRET experiments with each biosensor (white circles, control; grey squares, LPS; grey circle, Thiamet G; black squares, LPS+ Thiamet G). Right panels correspond to the increase BRET above basal induced by LPS, Thiamet G or LPS+Thiamet G (delta BRET expressed in milliBRET Units) and are the mean±SEM of 8 independent BRET experiments. Statistical analysis was performed using ANOVA followed by Dunnett’s post-test (*, **: p<0.05, p<0.01, respectively, when compared to the control condition). (B) RAW264.7 cells, bone marrow-derived and peritoneal primary mouse macrophages, and human monocyte-derived macrophages were cultured during 24h in absence or presence of LPS (100ng/mL), Thiamet G (10µM) or both. Proteins were extracted and analysed by western-blotting using an anti-O-GlcNAc antibody (RL2). Membranes were then re-probed with anti-clathrin antibody to control for protein loading in each well. Each blot is representative of at least 4 experiments.

RAW264.7 cells transfected with these biosensors were incubated for 24h in presence of LPS, Thiamet G (an inhibitor of OGA) or both. We observed that LPS treatment increased BRET signal with all three biosensors (**Fig. 1A**), indicating that TLR4 stimulation promotes a general rather than compartment-specific increase in O-GlcNAcylation in RAW264.7 macrophages. Interestingly, the effects of LPS and Thiamet G were additive, suggesting that the effect of LPS on O-GlcNAcylation is independent of OGA activity.

LPS-induced O-GlcNAcylation of proteins, both in absence and presence of Thiamet G, was further demonstrated by western-blotting using anti-O-GlcNAc antibody in RAW 264.7 cells, but also in mouse bone marrow-derived and peritoneal primary macrophages, as well as in human monocyte-derived macrophages (**Fig. 1B**). Therefore, increased O-GlcNAcylation upon LPS stimulation also appears to occur in primary macrophages from different origins and species, suggesting a general mechanism elicited by TLR4 activation.

### *GFAT2* is a TLR4-inducible gene in macrophages

We further explored, using RAW264.7 macrophages, the mechanism involved in this LPS-induced O-GlcNAcylation. As shown in **Figure 2**, LPS-induced increase in protein O-GlcNAcylation (**Fig. 2A**) was not associated with any detectable change in OGT and OGA mRNA or protein expression **(Fig. 2B and C)**, suggesting that it was not mediated by regulation of the expression level of O-GlcNAc cycling enzymes. GFAT, the enzyme that catalyzes the rate-limiting step of the hexosamine biosynthesis pathway **(Suppl Fig.1)**, exists as two isoforms (GFAT1 and GFAT2), encoded by two separate genes (also denominated *GFPT1* and *GFPT2*, respectively) that are differentially expressed in a cell-type specific manner. We observed that in the basal state, only GFAT1 protein was highly expressed in macrophages, whereas GFAT2 protein expression was barely detectable (**Fig. 2B**). GFAT1 protein expression was moderately increased (1.3 fold) by LPS treatment, whereas LPS induced a major increase (7 fold) in GFAT2 protein expression (**Fig. 2B**). In agreement with these results, **Figure 2C** shows that LPS increased the expression of GFAT1 mRNA by 2 to 3-fold, and markedly induced (by 10 to 15-fold) the expression of GFAT2 mRNA, suggesting a regulation of GFAT1 and 2 at the transcriptional level. Enzymatic assays indicated that whereas LPS treatment had no detectable effect on OGT or OGA activities, it significantly increased the activity of GFAT in macrophages (**Fig. 2D**). These results suggest that LPS-induced increase in the expression of GFAT, and more specifically GFAT2, translates into an increase in the activity of the rate limiting step of the hexosamine biosynthesis pathway. Time-course experiments indicated that a major induction of GFAT2 mRNA could be detected 1h after LPS stimulation, whereas GFAT1 mRNA expression was barely modified at early time points (**Suppl. Fig. S2**), revealing that *GFAT2* is an early TLR4-target gene.

**Figure 2:**
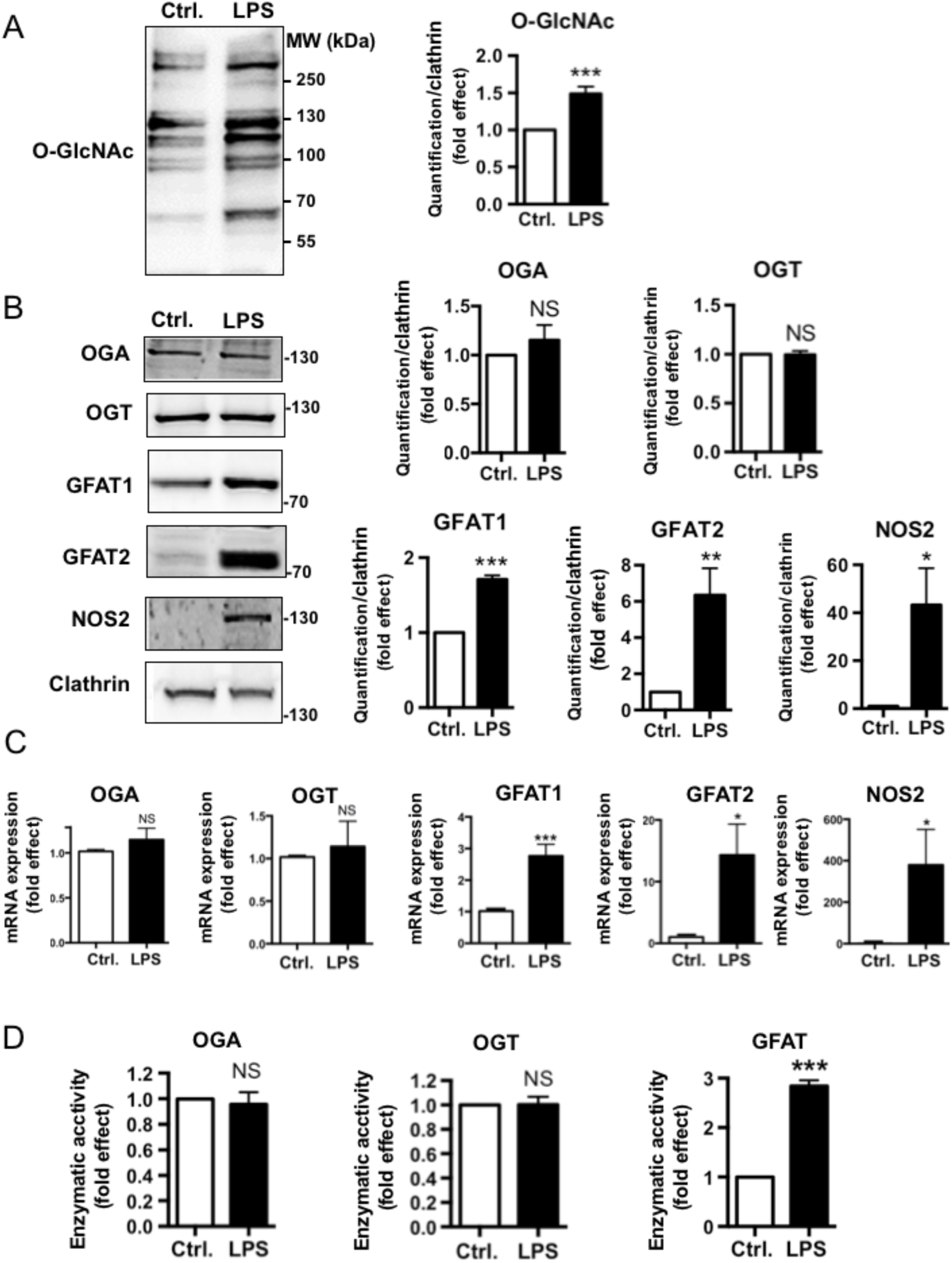
Effect of LPS on OGT, OGA and GFAT expression and activity in RAW264.7 cells. RAW264.7 cells were cultured during 24h in absence or presence of LPS (100ng/mL) and lysed for protein and RNA extraction. (A) O-GlcNAcylated proteins were precipitated on wheat-germ lectin (WGL) agarose beads and submitted to western-blotting using anti-O-GlcNAc antibody. Left panel, a typical western blot is shown. Right panel, the effect of LPS on global protein O-GlcNAcylation was quantified by densitometric analysis of the blots (B) Protein expression level of OGA, OGT, GFAT1, GFAT2, and NOS2 in total cell lysates. Left panel: a typical western blot is shown. Right panel, densitometric analysis of the blots. (C) mRNA expression levels of OGA, OGT, GFAT1, GFAT2 and NOS2 were evaluated by RT-qPCR. (D) OGT, OGA, and GFAT enzymatic activities in cell lysates from control and LPS-treated cells. Results are expressed as LPS-induced fold-effect and are the mean ± SEM of 4 to 8 independent experiments. Statistical analysis was performed using Student’s t test (*, **, ***: p<0.05, p<0.01, p<0.001, respectively; NS: non-significant).

Increased GFAT2 protein and mRNA expression was also observed in human monocyte-derived macrophages **(Fig. 3A)**, mouse bone marrow-derived macrophages **(Fig. 3B)**, as well as well as peritoneal macrophages (**Fig. 3C**). In agreement with these results, GFAT enzymatic activity was also increased upon LPS stimulation in human and mouse primary macrophages (**Suppl. Fig. S3**). Moreover, intraperitoneal injection of LPS in mice also induced a modest increase in GFAT1 and a marked increase in GFAT2 protein expression in peritoneal cells, indicating that this response was also operative *in vivo* **(Suppl. Fig. S4**).

**Figure 3:**
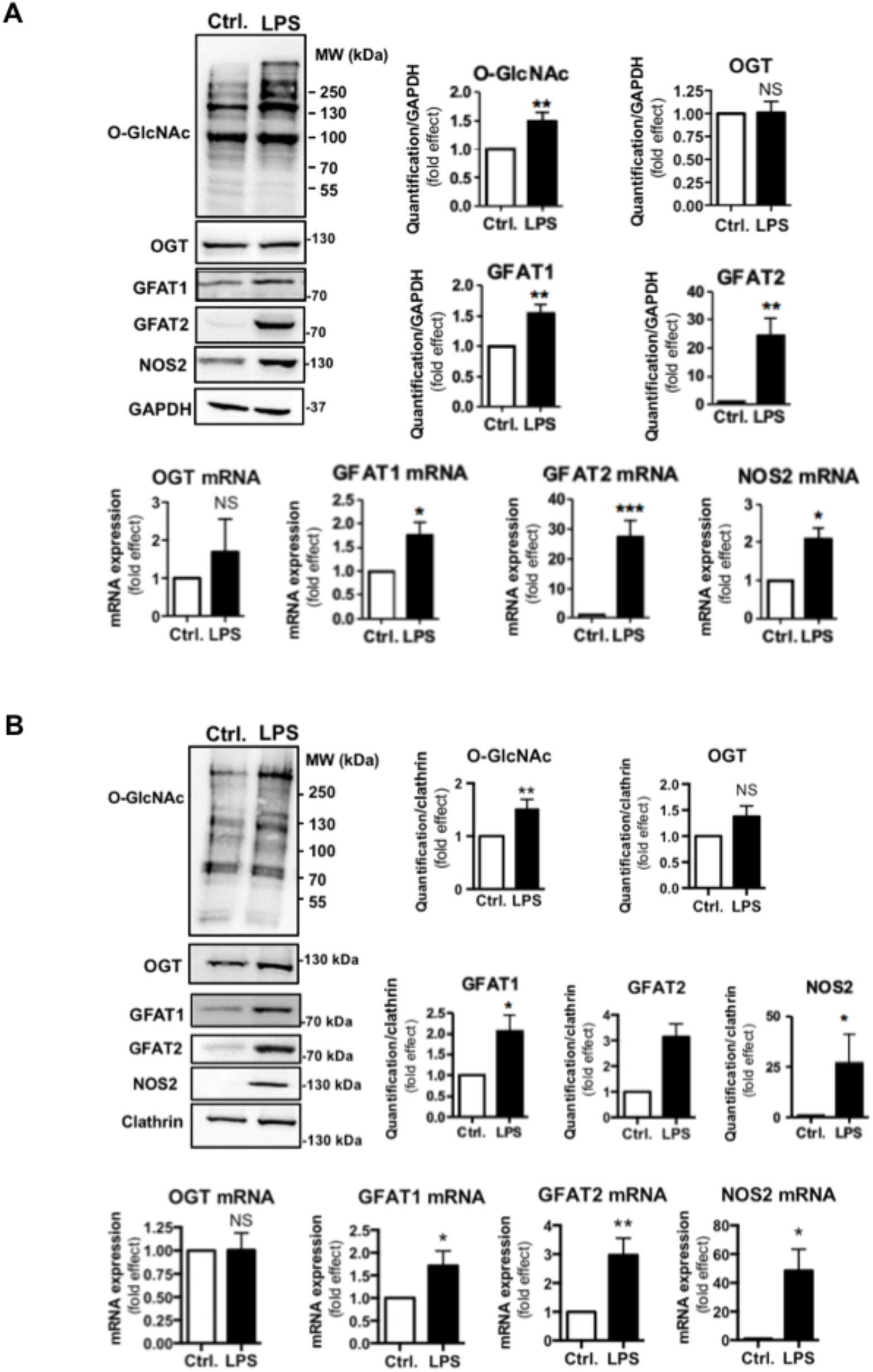
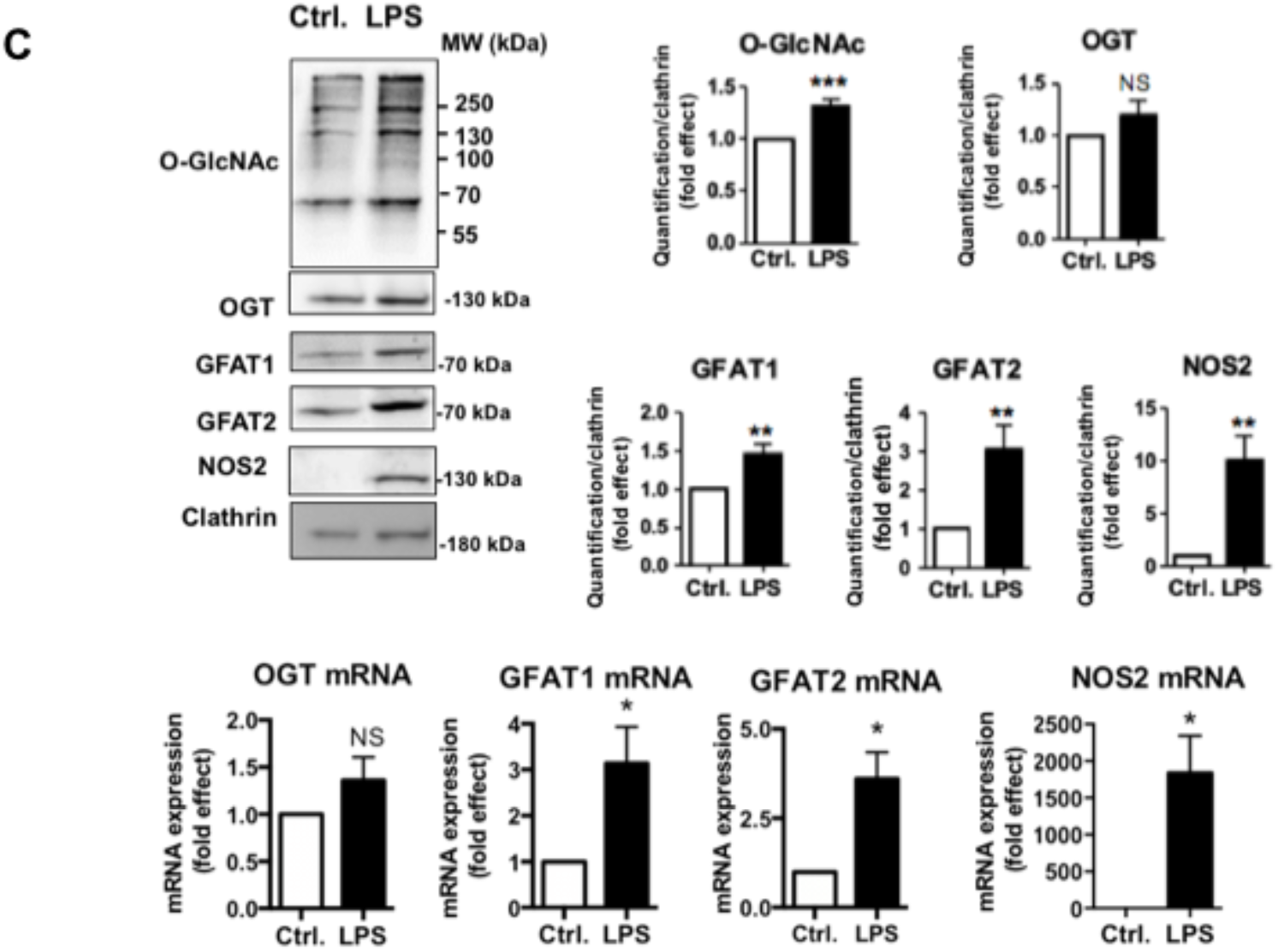
Effect of LPS on GFAT2 expression in human and mouse primary macrophages. Human monocyte-derived macrophages (A), mouse bone marrow-derived (B) and peritoneal (C) macrophages were cultured during 24h in absence or presence of LPS (100ng/mL) and lysed for protein and RNA extraction. The effect of LPS on global protein O-GlcNAcylation level and on the expression of OGT, OGA, GFAT1 and GFAT2 was evaluated by densitometric analysis of the western-blots. mRNA expression levels of GFAT2, GFAT1 and OGT were evaluated by RT-qPCR. Results are expressed as LPS-induced fold-effect and are the mean ± SEM of at least 4 independent experiments. Statistical analysis was performed using Student’s t test (*, **, *** p<0.05, p<0.01, p<0.001, respectively; NS: non-significant).

Altogether, our results suggest that LPS induces an increase in protein O-GlcNAcylation through stimulation of the expression of the rate-limiting enzyme of the hexosamine biosynthesis pathway.

### Impaired O-GlcNAcylation promotes pro-inflammatory response in macrophages

To evaluate the role of LPS-induced O-GlcNAcylation in macrophages, we isolated peritoneal macrophages from mice with conditional deletion of OGT in the myeloid lineage. As expected, OGT expression and LPS-induced O-GlcNAcylation were markedly impaired in OGT-KO cells (**Fig. 4A**). We noticed that basal GFAT1 and both basal and LPS-induced GFAT2 expression were increased in OGT-KO macrophages, suggesting a compensatory response to the low level of O-GlcNAcylation in these cells. Interestingly, OGT deletion resulted in an increased in LPS-induced NOS2 expression. Moreover, we observed an increased production of IFNγ and IL1β in the culture medium of OGT-KO macrophages. These results suggest that increased O-GlcNAcylation upon LPS stimulation may have a counter-regulatory effect that restrains excessive cytokines production by macrophages (**Fig. 4B**). In agreement with this notion, significant increases in pro-inflammatory cytokines IFN γ, IL1β and TNFα were also observed *in vivo* in the serum of OGT-KO mice intraperitoneally injected with LPS (**Fig. 4C**).

**Figure 4:**
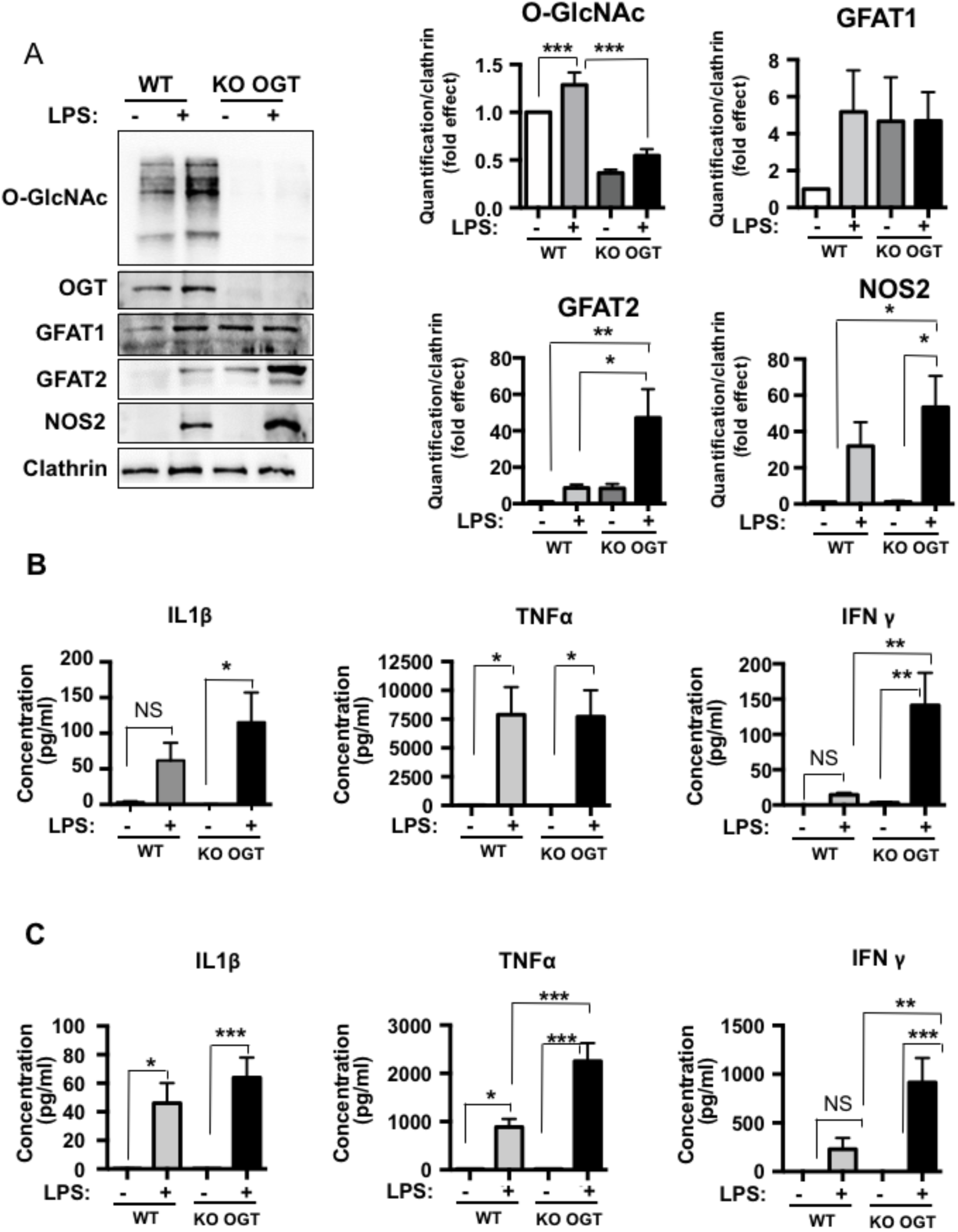
Targeted deletion of OGT in macrophages increases the pro-inflammatory effect of LPS. (A, B) Peritoneal macrophages from control or OGT-KO mice were cultured during 24h in absence or presence of LPS (100ng/mL). The culture medium was collected for measurement of cytokine production in the medium, and cell were lysed for protein analysis. (A) Typical western blot showing protein O-GlcNAcylation and expression of OGT, GFAT1, GFAT2 and NOS2 in control and OGT-KO macrophages. (B) The concentration of IL1β, TNFα and IFNγ secreted in the culture medium by control and OGT-KO macrophages were determined using MSD kit. (C) Control and OGT-KO mice were intraperitoneally injected with 0.6 mg/kg of LPS. 6h after injection, mice were sacrificed and blood was collected. The concentration of IL1β, TNFα and IFNγ in the serum were determined using MSD kit. Results are expressed are the mean ± SEM of 4 to 9 independent experiments. Statistical analysis was performed using ANOVA followed by Tukey’s post-test (*, **, ***: p<0.05, p<0.01, p<0.001, respectively).

These results suggest that increased O-GlcNAcylation upon TLR4 activation participates in the counter-regulation of proinflammatory signalling in macrophages. Induction of GFAT2 expression may therefore constitute a new mechanism to limit excessive inflammation upon LPS stimulation.

### GFAT2 expression is dependent on FoxO1 transcription factor in macrophages

The transcription factor FoxO1 have been shown previously to display dual regulatory effects on inflammatory signals in myeloid cells, with either pro- [49-52] or anti-inflammatory [53-56] effects, depending on the pathophysiological context. Analysis of mouse and human GFAT2 putative promoters revealed canonical FoxO1 recognition sites within the 500bp and 800pb regions upstream of the transcription start site of the mouse and human genes, respectively (**Suppl. Fig. S5**). We inserted these upstream sequences in a luciferase reporter gene plasmid. Transfection in HEK293 cells with this plasmid revealed that the activities of both human and mouse reporter genes were markedly increased by co-transfection with a constitutively active form of FOXO1, FOXO1-TM (**Fig. 5A)**.

**Figure 5:**
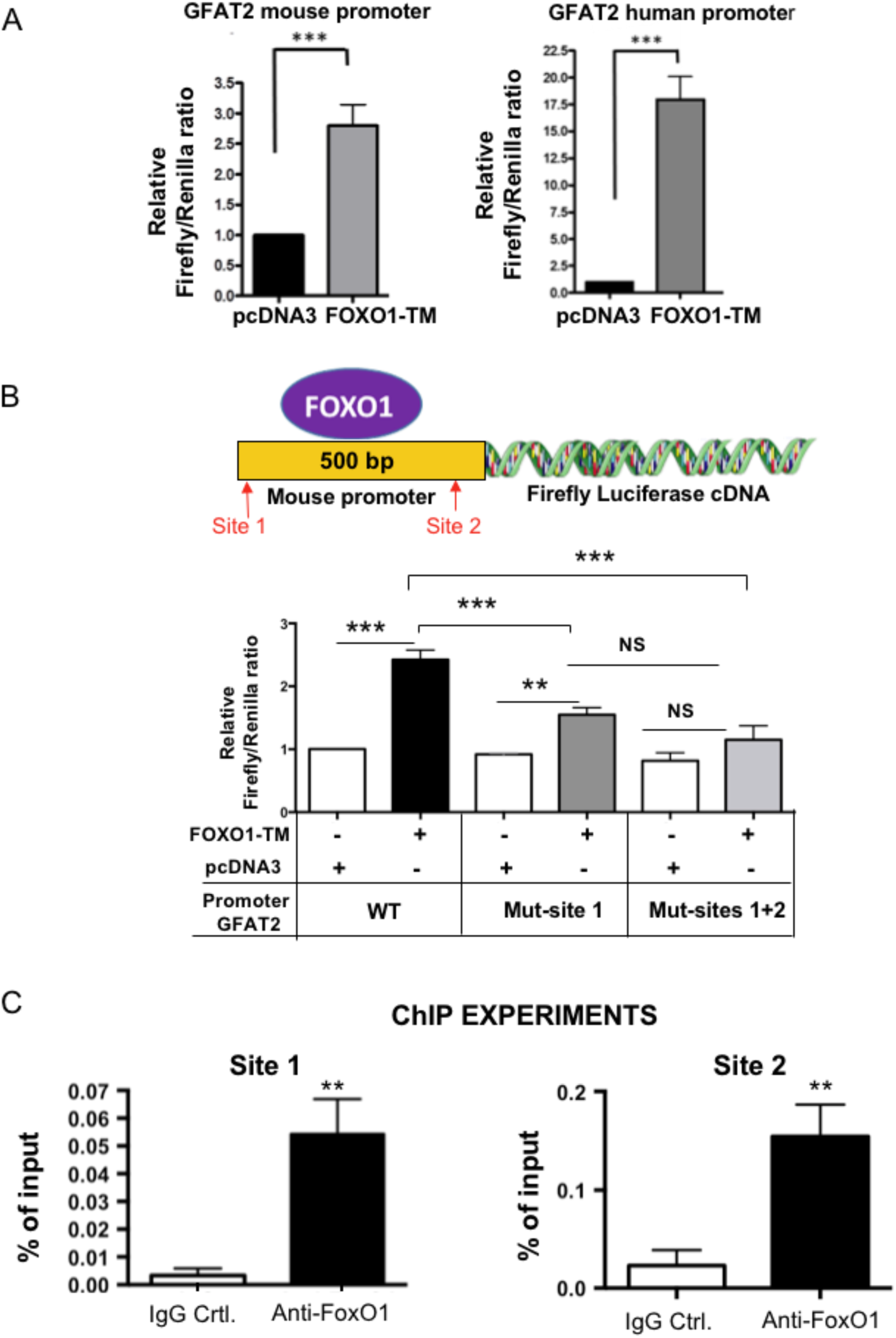
FoxO1 stimulates the activity of a GFAT2 promoter reporter gene. (A) HEK 293-T cells were co-transfected with a firefly luciferase reporter gene comprising the 500bp or 800pb regions upstream of the transcription start site of the mouse and human genes, respectively, a *Renilla* luciferase expression plasmid, and either pcDNA3 or the cDNA coding for the constitutively active FOXO1-TM. Results are expressed as fold-activation of the promoter by FOXO1 and are the mean ± SEM of 5 independent experiments. Statistical analysis was performed using Student’s t test (***: p<0.001). (B) Mutation of FoxO1 binding sites on the mouse GFAT2 promoter inhibits FoxO1 induced activation of the GFAT2 promoter reporter gene. Results are expressed as fold-effect of the basal activity of the wild-type promoter and are the mean ± SEM of at least 5 independent experiments. Statistical analysis was performed using ANOVA followed by Tukey’s test (**, ***: p<0.01, p<0.001, respectively; NS: non-significant). (C) Binding of Foxo1 on Site 1 and Site 2 of the endogenous mouse promoter was shown by chromatin immunoprecipitation of Foxo1 in RAW264.7 cells. Chromatin was immunoprecipitated with either control IgG or anti-FoxO1 antibody and recovered DNA was amplified by qPCR. Results are the mean ± SEM of 5 independent experiments, each performed in biological duplicate. Statistical analysis was performed using Student’s t test (**, p<0.01).

The mouse 500bp promoter contains two FoxO1 recognition sites (referred to as Site1 and Site 2 on **Fig. 5B** and **Suppl Fig. S5**). To further demonstrate the contribution of these sites, we mutated either one or both FoxO1 binding sites and measured the activity of the mutated promoters in the luciferase assay.

As shown in **Fig. 5B**, FOXO1-TM effect was markedly reduced by mutation of Site 1 and totally abolished when both Site 1 and Site 2 were mutated. This suggests that both sites might be important for regulation of GFAT2 expression by FoxO1. Binding of Foxo1 to Site 1 and Site 2 was further demonstrated on the endogenous GFAT2 promoter in RAW264.7 cells, in chromatin immunoprecipitation experiments using a FoxO1-specific antibody (**Fig. 5C).**

These results suggested that FoxO1 may be involved in the regulation of GFAT2 expression in macrophages. To evaluate the potential involvement of FoxO1 in the regulation of GFAT2 expression, we used a small molecule specific inhibitor of FoxO1 activity, the AS1842856 drug [57]. We observed, both in RAW264.7 cells (**Fig. 6A**) and human monocyte-derived macrophages (**Fig. 6B**), that inhibition of FoxO1 by this compound markedly impaired the effect of LPS on induction of GFAT2 mRNA and protein expression.

**Figure 6:**
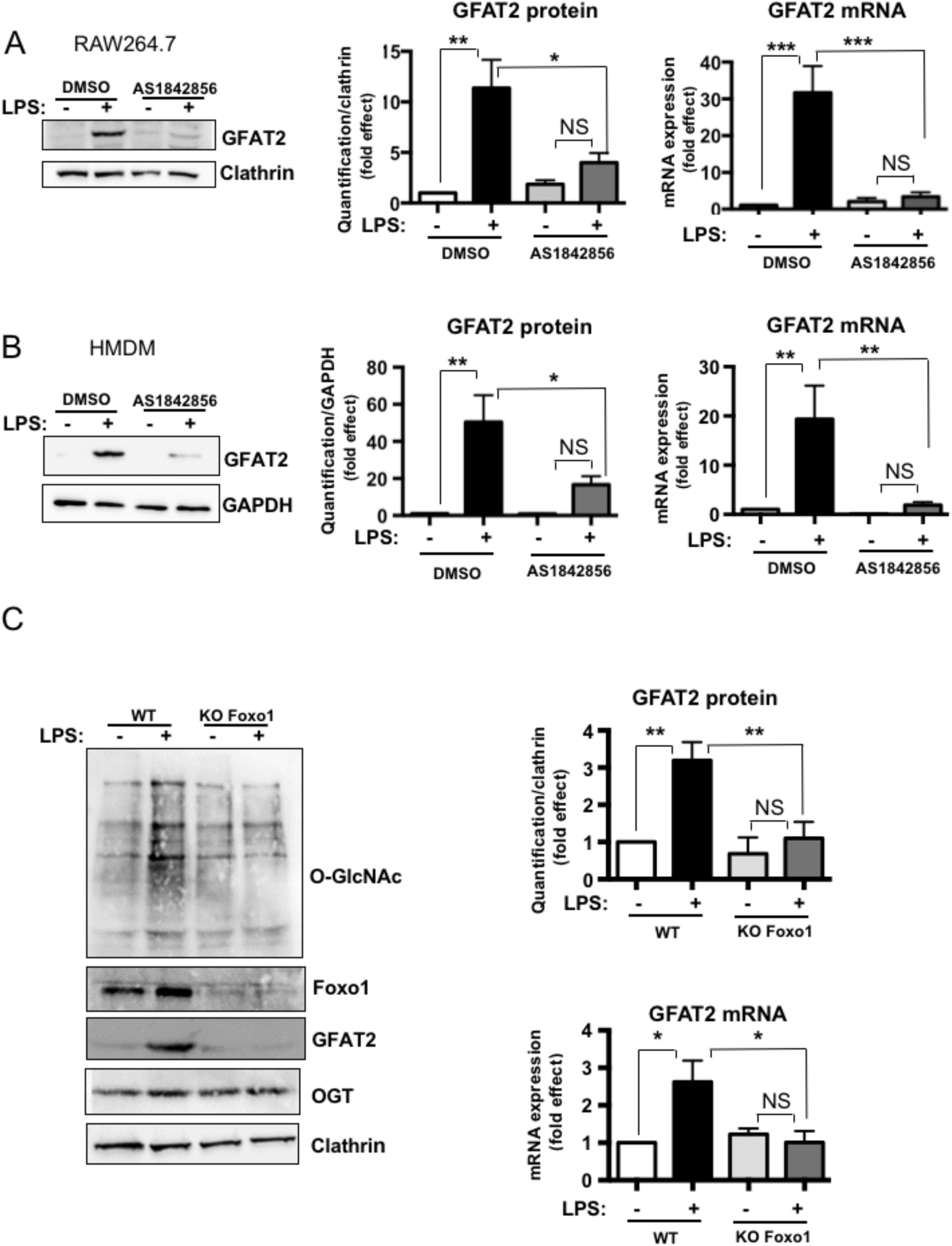
Pharmacological or genetic inhibition of FoxO1 impairs LPS effect on GFAT2 mRNA and protein expression. RAW 264.7 cells (A) and human monocyte-derived macrophages (B) were cultured for 3 days in the presence of the FoxO1 inhibitor AS1842856 (500nM) or vehicle only and then incubated for an additional 24h in absence or presence of LPS (100ng/mL) and lysed for protein and RNA extraction. Left panels: typical western blots showing the effect of LPS on GFAT2 protein expression. Right panels, quantification of the western-blot and expression of GFAT2 mRNA evaluated by RT-qPCR. Results are expressed as fold-effect of control condition and are the mean ± SEM of 5 to 6 independent experiments. Statistical analysis was performed using ANOVA followed by Tukey’s test (***, p<0.001, NS: non-significant). (C) Effect of LPS on protein O-GlcNAcylation and GFAT2 protein and mRNA expression in bone-marrow derived macrophages from wild-type and Foxo1-KO mice. Results are expressed as fold-effect of wild-type control condition and are the mean ± SEM of 3 to 5 independent experiments. Statistical analysis was performed using ANOVA followed by Tukey’s test (*, **, ***: p<0.05, p<0.01, and p<0.001, respectively; NS: non-significant).

To further demonstrate the role of Foxo1 in LPS-induced GFAT2 expression, we also evaluated the effect of LPS on peritoneal macrophages from mice with conditional Foxo1 deletion in the myeloid lineage. We observed that LPS-induced GFAT2 protein and mRNA expression was markedly impaired in Foxo1 KO macrophages (**Fig. 6C).** Inhibition of GFAT2 expression was associated with inhibition of LPS effect on protein O-GlcNAcylation. These results demonstrate that Foxo1 is necessary for induction of GFAT2 by LPS and support the notion that increased GFAT2 mediates LPS-induced increase in protein O-GlcNAcylation in macrophages.

## Discussion

Several lines of evidence have suggested a role for O-GlcNAcylation in the regulation of inflammatory processes in macrophages [58]. However, contradictory results have been obtained concerning the effect of TLR4 activation on protein O-GLNAcylation. Indeed, depending on the experimental setting, both increases [22,23] and decreases [24,25] in the general O-GlcNAcylation profile were observed upon LPS stimulation.

In the present work, we showed, using two different methodological approaches (BRET-based assay (**Fig. 1A**) and western-blotting on crude cell lysate as well as WGL-bound fraction (**Fig. 1B and 2A**)) that LPS induced a major increase in global protein O-GlcNAcylation in RAW264.7 macrophages. The use of plasma membrane-, cytosol- or nucleus-targeted BRET O-GlcNAc biosensors indicated that the LPS-induced increased in O-GlcNAcylation was not restricted to a specific cell compartment. Increased protein O-GlcNAcylation upon LPS treatment was also confirmed in primary mice and human macrophages, indicating that our observation was not a cell line specific effect. The reason underlying the discrepancies between different laboratories are unknown at the present time. However, it is possible technical differences in the extraction procedure may affect the detection of O-GlcNAc by western-blotting. For instance, whereas investigators generally add proteases and phosphatases inhibitors in their cell lysis buffer, they do not mention the use of any hexosaminidase inhibitor to prevent loss of O-GlcNAc during the extraction procedure. Given that O-GlcNAc is a very dynamic and labile modification, that can be rapidly hydrolyzed upon cellular damage or during protein isolation [59], it is quite possible that loss of O-GlcNAc might occur during the sample preparation. In contrast, we always included PUGNAc at a concentration of 10μM in our extraction buffer, in order to preserve O-GlcNAcylation state of proteins obtained in cells after LPS treatment. In addition, the confirmation of our western-blotting results by an independent technique based on the use of a BRET biosensor, which monitors O-GlcNAcylation changes in intact living cells without any processing of cellular proteins, strongly argues in in favour of an LPS-induced general increase in O-GlcNAcylation in macrophages.

Interestingly, the effect of LPS on protein O-GlcNAcylation were additive to those of a maximally inhibitory concentration of the OGA inhibitor, Thiamet G. This suggested that LPS-induced O-GlcNAcylation was not mediated by regulation of OGA activity. In agreement with this notion, using a fluorogenic substrate, we observed that OGA activity was similar in cell extracts from control and LPS stimulated RAW264.7 macrophages (**Fig.2 D**). Moreover, using a luminescent assay [33,42], we found that LPS treatment had no detectable effect on OGT activity in cell extracts from RAW264.7 cells (**Fig. 2D**). Therefore, LPS-induced O-GlcNAcylation does not appear to be mediated by regulation of O-GlcNAc cycling enzymes.

One of the most important finding of our study is that LPS treatment resulted in an increased expression and activity of GFAT **(Fig. 2)**. GFAT is the enzyme that catalyses the rate limiting step of the hexosamine biosynthesis pathway, which eventually leads to the production of UDP-GlcNAc, the substrate used by OGT for protein O-GlcNAcylation. GFAT exists in two isoforms, GFAT1 and GFAT2, encoded by two different genes [60,61]. Although little data are available concerning differential roles of these enzymes, these two isoforms present different tissue distribution, with GFAT1 mRNA being predominantly expressed in pancreas, placenta and testis, whereas GFAT2 mRNA were found throughout the central nervous system [61]. Whereas some differences in the regulation of their catalytic activities by cAMP-induced phosphorylation has been described [62], very little is known about differential regulation of GFAT1 versus GFAT2 expression in different cell types. Most surprisingly, to our knowledge, only one study evaluated in macrophages the expression of GFAT in a mouse macrophage cell line (ANA-1) [63]. These authors reported that GFAT1 was constitutively expressed in these cells and they indicated (as data not shown) that no effect of LPS or IFNγ on its expression was observed. However, GFAT2 expression was not evaluated in this study. Interestingly, we found that GFAT1 mRNA and protein were indeed expressed at significant levels in resting macrophages, whereas GFAT2 expression was barely detectable. LPS stimulated both GFAT1 and GFAT2 expression, although the stimulatory effect was much higher for GFAT2, which turned-out to be an early TLR4-inducible gene in macrophages (**Suppl. Fig 3**). Thus, LPS-induced increase in O-GlcNAcylation is likely to be at least in part mediated by the induction of the expression of the GFAT2 isoform in macrophages.

Although the exact mechanism by which TLR4 stimulates GFAT2 expression remains elusive, we provide strong evidence for a role of FoxO1 in this process. Indeed, using gene reporter as well as chromatin immunoprecipitation assays, we demonstrated the presence of FoxO1 binding sites on GFAT2 putative promoter. Moreover, pharmacological inhibition or genetic deletion of FoxO1 in macrophages markedly impaired LPS-induced GFAT2 expression, confirming the involvement of FoxO1 in this regulation.

Previous studies have shown that O-GlcNAcylation can either promote inflammation or reduces it, according to the cellular context and type of insult. Thus, whereas O-GlcNAcylation has pro-inflammatory effects in situations of chronic hyperglycaemia, it appears to be protective in acute stress conditions, such as ischemia-reperfusion injury in the heart [58]. We observed that impaired O-GlcNAcylation in OGT-KO macrophages resulted in marked increase in NOS2 expression and pro-inflammatory cytokines production, suggesting that the O-GlcNAc tone may exerts a break on inflammatory processes. Therefore, the rapid induction of GFAT2 expression may constitute a protective mechanism to limit exacerbated inflammation upon LPS stimulation.

In summary, we have shown that LPS stimulation promotes a general increase in protein O-GlcNAcylation in macrophages. This effect is at least in part mediated by increased expression and activity of the rate-limiting enzyme of the hexosamine biosynthesis pathway, GFAT, with the GFAT2 isoform being the most responsive to TLR4 activation. Indeed, while GFAT1 may control the activity of the HBP in the basal state, our work revealed that *GFAT2* is a new TLR4-inducible gene in macrophages, permitting a rapid adaptive response to environmental changes.

## Acknowledgments

We are very grateful to Prof. L.K. Mahal for the cDNA coding for the FRET O-GlcNAc biosensors. We also thank Laura Francese for her help in some of the BMDM experiments. This work was performed within the Département Hospitalo-Universitaire AUToimmune and HORmonal diseases.

