## Supplemental Data for "LPS induces GFAT2 expression to promote O-GlcNAcylation and attenuate inflammation in macrophages"

Running title: **GFAT2 is a FoxO1-dependent TLR4-inducible gene**

Hasanain AL-MUKH, Léa BAUDOIN, Abdelouahab BOUABOUD, José-Luis SANCHEZ-SALGADO, Nabih MARAQA, Mostafa KHAIR, Patrick PAGESY, Georges BISMUTH, Florence NIEDERGANG and Tarik ISSAD

*Université de Paris, Institut Cochin, CNRS, INSERM, F-75014 Paris, France*

Address correspondence to: Tarik Issad, Institut Cochin, Department of Endocrinology, Metabolism and Diabetes, 24 rue du Faubourg Saint-Jacques, 75014 Paris FRANCE. Tel : + 33 1 44 41 25 67;

### Supplementary Figures

#### Supplementary Fig. S1

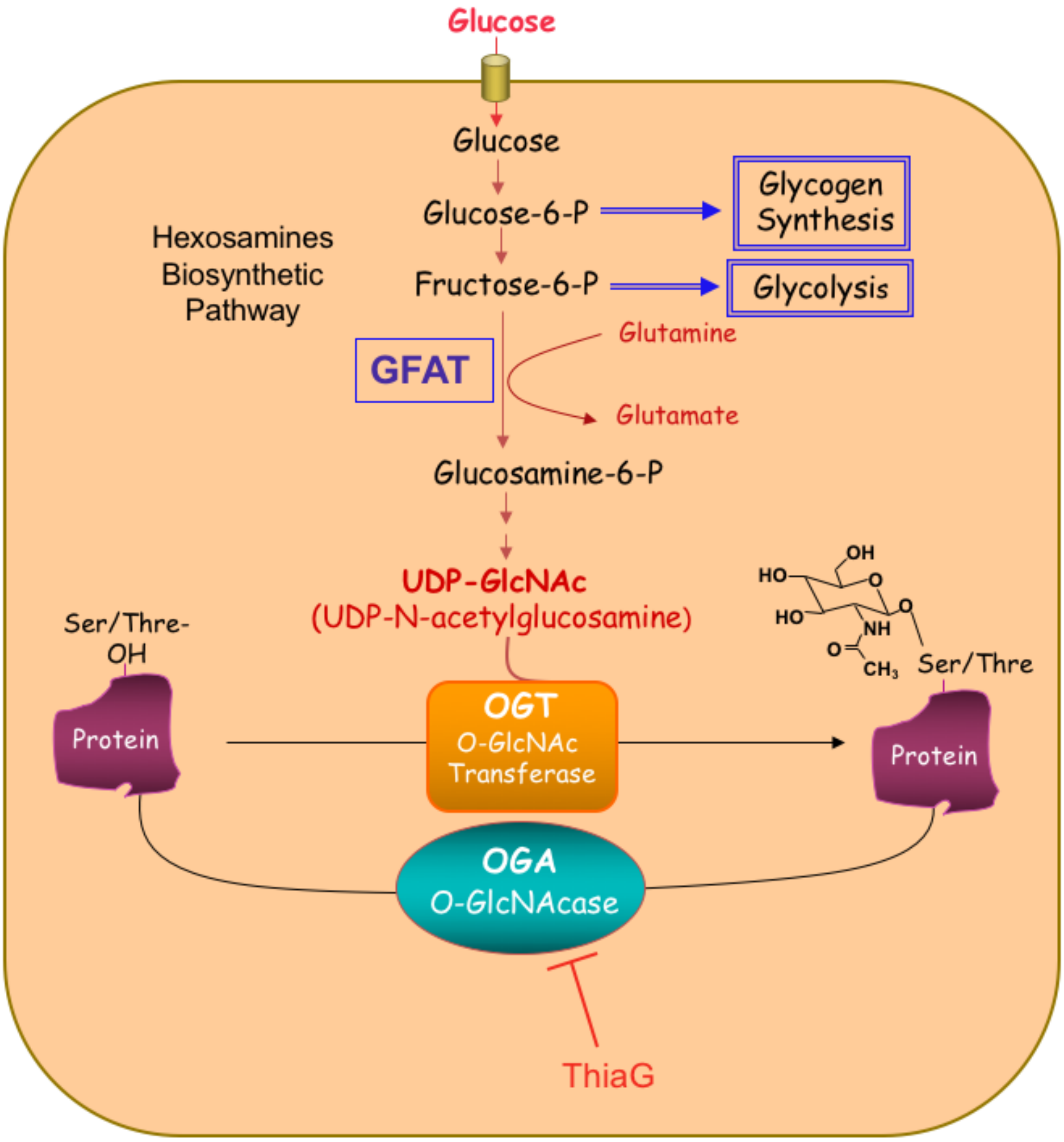

**Supplementary Fig. S2**

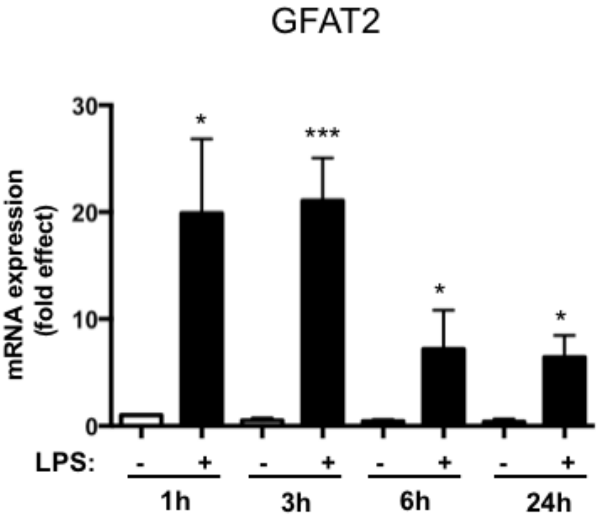


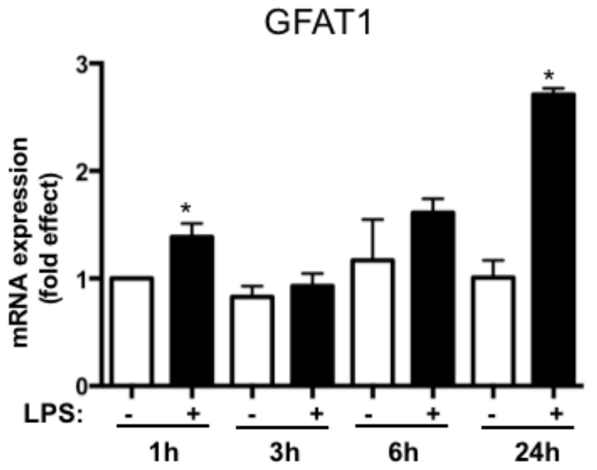

### Supplementary Figure S3

**GFAT activity**

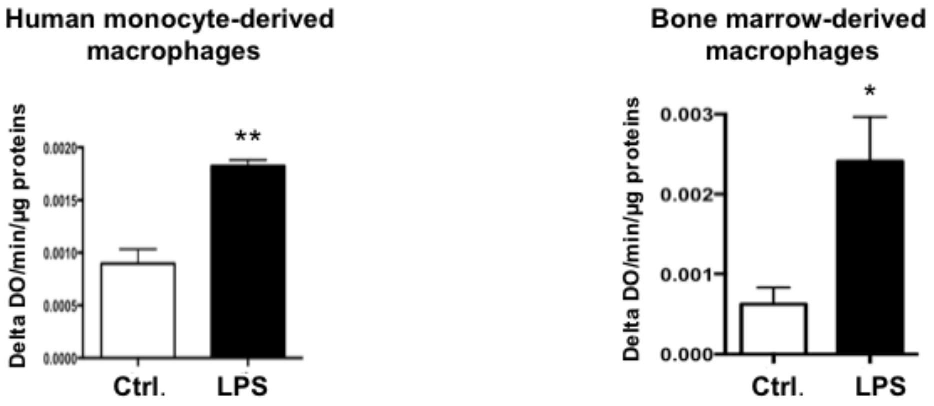

### Supplementary Fig. S4

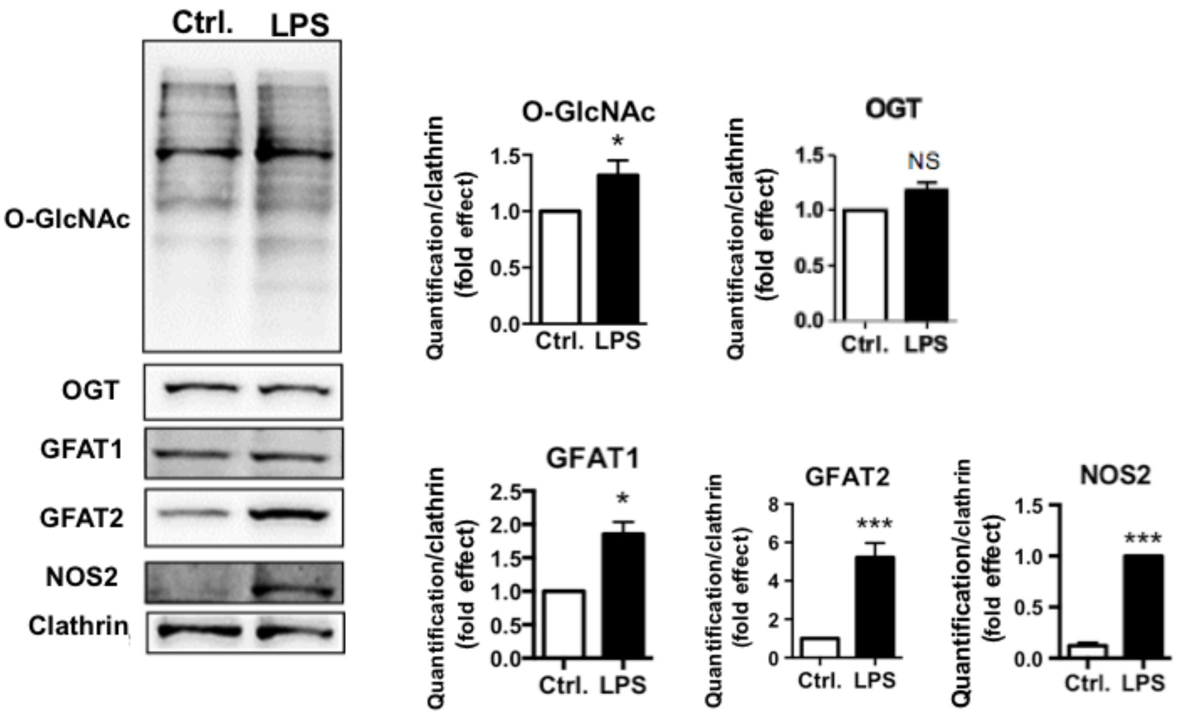

### Supplemeary Fig. S5

A

**Human GFAT2 promoter: -801>>0**

TCAGGCAATCCACCCATCTTGGCCTCCCAAAGTGCTGGGATTACAGGCATGAGCCACCAC
GCAGTGCCTGAACCACTGCACCCGGCCAATTTTTTTCTTACAGAAGAAATACATGTTCAT
TTT**AGGAAACAGAGA**ATAAAAATATTTTTAAAGGGTTAATAACTCATCTTTTTACCTACC
ACCAAATGTTATTCTCTGCATCTATAGTCAGTATTTAAACAACAATACAGTCATTCTGCA
AACATTGTTTTGTAGCTGGAATTTGACTTCTGGTTATAGAGAACTTCAAATTGTGCTGCA
AAGCTGGGATCTGGCCCAGAGTCGTCAGCGCTGTCCCTGATGCACCTCTGTGCCACGGCA
TCCTGGAAAGCTTGGGCTTTCCACTCTGCTTGGCCAGGAAAGGGTGGGGGTTTGTGCGAG
ACAAGCACTAATCTGATATTGCACTAAGCATTGACTGAGAGACGGCGCTGGTGGAGGGC
AGCAGGCTACCCCGTCGCCCTGTTCCTGTTAACTCTCTGGCAGAGACAAGGGAGCTGGGG
CTGTGGGCTGAATGTTTGTGTCCCCCCAAATTTATATGTTGAAATCCTAGCCCCCAAGAT
GATGCTATTAGGATACAGGGCCTTTTGGGAGGTGACTCAGTCATGATGACTGAGGCCTCA
TGACGGGGTTAGTGCCTTACCTGAGGGACCCCAGAGAGCTACCTTGGCCCCTTGCGATAC
AGTGAGAAAGCACCGTCCATGAACCAGGAAATGAGCCCTCAGGAGACAGCAAATCTGCTG
GTGCCTTGATCTTGGACTTCC

B

**Mouse GFAT2 promoter: -500>>0**

GGCAAGTGCTCTACTACTGAGCTAAGTCCTTGATCCTGGCTCTCTTTATAAAGTGGCC
TGCCTCTGATATTTCAT**GTAAACACAC**ACACACACACACACACACAGAGAGAGAGAGA
GAGAGAGAGAGAGAGAGAGAGAGAGAGAAACACAGAGAGAGACAGAGAGACAGACAAT
ACTTGTCCCCATTGGTTTATGTCCAGTTGTTTTCCTCTACAGAACACATTGGATTCTTCA
GTTAGATTCCTCAGCAGTGGTAGCAGTAGGGGAGAAGAGAAGAAATGTTTGAAAAGAGTA
GAAACGGAGAACAGGGAGACAGAGCAGCATCCCCAACAGGCAAGAATTGCTGTCTTTACT
ACTTCCTACAGAATTAGGCCCAGAGGAGCTAATCCCTTGCCCAAGGGAGTATTGCTGATA
AGTAGAATTCAATTG**GAACACAGAC**CAAGGAAGAGCTTGCATAGAAGCAGGCCTCTGACA
AATTCTTGTTATCCTAGCACCC

GTAAACACAC Site 1
GAACACAGAC Site 2

**Supplementary Figure legends**

**S1: The hexosamine biosynthetic pathway and protein O-GlcNAcylation**

The hexosamine biosynthetic pathway (HBP) flux controls O-GlcNAcylation of intracellular proteins. This dynamic and reversible post-translational modification controls the activity, the localization and/or the stability of proteins, according to the rate of glucose entering the HBP. Fructose-6-phosphate is converted to glucosamine-6-phosphate by the glutamine:fructose-6-phosphate amidotransferase (GFAT), the rate limiting enzyme of the pathway. After a subset of reactions, UDP-N-acetylglucosamine (UDP-GlcNAc) is generated and used by the O-GlcNAc transferase (OGT) as a substrate to add GlcNAc on serine or threonine residues of target proteins. O-GlcNAc moiety is removed from O-GlcNAc-modified proteins by the O-GlcNAcase (OGA). ThiaG: Thiamet G, a highly selective inhibitor of OGA.

**S2: Time-course experiment of GFAT2 induction by LPS in RAW264.7 cells.**

RAW264.7 cells were cultured in absence or presence of LPS (100ng/mL) for 1, 3, 6, and 24h. RNA were extracted and GFAT2 expression was analysed by RT-qPCR. Results are expressed as fold-effect relative to basal level at 1h, and are the mean±SEM of at least 3 independent experiments. Statistical analysis was performed using Student's t test (*, ***: $p<0.05$ and $p<0.001$, respectively).

**S3: Effect of LPS on GFAT enzymatic activity in human and mouse primary macrophages.**

Human monocyte-derived and mouse bone-marrow-derived macrophages were cultured during 24h in absence or presence of LPS (100ng/mL) and lysed for protein extraction. GFAT enzymatic activity was measured as described in the method section.

Results are the mean±SEM of 3 to 5 experiments. Statistical analysis was performed using Student's t test (*, **: $p<0.05$ and $p<0.01$, respectively).

**S4: Effect of *in vivo* administration of LPS on O-GlcNAcylation profile and GFAT2 protein expression in peritoneal cells.** Mice were injected intraperitoneally with 0.6 mg/kg of LPS. 6h after injection, mice were sacrificed and peritoneal cells were collected. Left panel: A typical western blot showing the effect of LPS on global protein O-GlcNAcylation level and expression of OGT, OGA, GFAT1, GFAT2 and NOS2 is shown. Right panel: Densitometric analysis of the blots. Results are expressed as fold-effect and are the mean ±

SEM of at least 4 independent experiments. Statistical analysis was performed using Student's t test (*,***: $p<0.05$, $p<0.001$, respectively; NS: non-significant).

**S5: FoxO1 binding sequences in the GFAT2 promoter.** (A) Human (-801>>>0 ) and (B) mouse GFAT2 (-501>>>0) putative promoter sequences used for luciferase assays. FoxO1 binding sites are highlighted in green and blue.

**Supplementary Table 1: List of antibodies**

| | |
|---|---|
| Anti GFAT2 antibody | Abcam (EPR 19095) ab190966 |
| Anti-GFAT1 antibody | Santa Cruz Biotechnology (H-49) : sc-134894 |
| Anti-FOXO1 antibody | Cell signaling(L27) Antibody 9454 |
| Anti-FOXO1A antibody - Chip Grade | Abcam (ab39670) |
| Anti-NOS2 polyclonal antibody | Santa Cruz Biotechnology (N-20) : Sc-651 |
| Anti-O-GlcNAc Transferase OGT antibody | Sigma (O 6264) |
| Anti-O-GlcNAcase/OGA/MGEA5 antibody | Novus bio (NBP2-32233) |
| Anti-O-GlcNAc (RL2) antibody | Thermo Fisher (MA1-072) |
| Anti-GAPDH monoclonal antibody | Santa Cruz Biotechnology (O411) sc-47724 |
| Anti-Clathrin antibody | BD transduction labratories (610500) |
| HRP-conjugated anti-rabbit antibody | Santa Cruz Biotechnology |
| HRP-conjugated anti-mouse antibody | Jackson ImmunoResearch, Laboratories |

**Supplementary Table 2:** Sequences of the primers

**Mouse primers used for RT-qPCR**

**Cyclophiline**
Forward: ATGGCACTGGCGGCAGGTCC
Reverse: TTGCCATTCCTGGACCCAAA
**HPRT**
Forward: GCTGGTGAAAAGGACCTCT
Reverse: CACAGGACTAGAACACCTGC
**NOS2**
Forward: CCAAGCCCTCACCTACTTCC
Reverse: CTCTGAGGGCTGACACAAGG
**OGT**
Forward: GCCCTGGGTCGCTTGGAAGA
Reverse: TGCCACAGCTCTGTCAAAAA
**OGA**
Forward: CGGGAATTCCAGTGGCTTCG
Reverse : AAGCCGGGTGAACATCCCCA
**GFAT1**
Forward: GAGACAGATTGCGGGGTTGA
Reverse: CGGCAGTCGCTTCAGTCC
**GFAT2** Forward: GTATGATTGGCCGACCCTGG
Reverse: ATGCTAGCCGGAGAGCTGAA
**GFAT2 promoter site 1 – CHIP**
Forward: AAGTGCTCTACTACTGAGCTAAGTCC
Reverse: GCTACCACTGCTGAGGAATCTAACTGA
**GFAT2 promoter site 2 – CHIP**
Forward: AGAGCAGCATCCCCAACAGGCA
Reverse: ATAACAAGAATTTGTCAGAGGCCTGC

**Human primers used for RT-qPCR**
**NOS2**
Forward: CAGCGGGATGACTTTCCAA
Reverse: AGGCAAGATTTGGACCTGCA
**GFAT1**
Forward: GGA TAT GAT TCT GCT GGT GTG
Reverse: CCA ACG GGT ATG AGC TAT TC
**GFAT2**
Forward: CGGAGTCCGGAGCAAATACA
Reverse: AAGATGACCCGGTTGGTGTG
**ARN18S**
Forward: GTAACCCGTTGAACCCCATT
Reverse: CCATCCAATCGGTAGTAGCG

**Mouse oligonucletorides for mutagenesis**
**GFAT2 promoter site 1 - mutagenesis**
Forward: GATATTTCATGTAACGACACACACACACACAC
Reverse: GTGTGTGTGTGTGTGTCGTTACATGAAATATC
**GFAT2 promoter site 2 - mutagenesis**
Forward: GATAAGTAGAATTCAATTGGAACCGAGACCAAGGAAGAGCTTGCATAG
Reverse: CTATGCAAGCTCTTCCTTGGTCTCGGTTCCAATTGAATTCTACTTATC
